# The NADPH oxidase NOX2 is a marker of adverse prognosis involved in chemoresistance of Acute Myeloid Leukemias

**DOI:** 10.1101/2021.03.04.433871

**Authors:** Paolillo Rosa, Boulanger Mathias, Gâtel Pierre, Gabellier Ludovic, Tempé Denis, Hallal Rawan, De Toledo Marion, Moreaux Jérome, Baik Hayeon, Elise Gueret, Récher Christian, Sarry Jean-Emmanuel, Cartron Guillaume, Piechaczyk Marc, Bossis Guillaume

## Abstract

Resistance to chemotherapeutic drugs is a major cause of treatment failure in Acute Myeloid Leukemias (AML). To better characterize the mechanisms of chemoresistance, we first identified genes whose expression is dysregulated in AML cells resistant to daunorubicin (DNR) or cytarabine (Ara-C), the main drugs used for the induction therapy. The genes found activated are mostly linked to immune signaling and inflammation. Among them, we identified a strong up-regulation of the NOX2 NAPDH oxidase subunit genes (*CYBB*, *CYBA*, *NCF1*, *NCF2*, *NCF4* and *RAC2*). The ensuing increase in NADPH oxidase activity, which is particularly strong in DNR-resistant cells, participates in the acquisition and/or maintenance of resistance to DNR. In addition, analyzing gp91^phox^ (*CYBB*-encoded Nox2 catalytic sub-unit) expression at the surface of leukemic blasts from 74 patients at diagnosis showed that NOX2 is generally more expressed and active in leukemic cells from the FAB M4/M5 subtypes compared to FAB M0-M2 ones. Using a gene expression-based score we demonstrate that high NOX2 subunit genes expression is a marker of adverse prognosis, independent of the cytogenetic-based risk classification, in AML patients.

## Introduction

Acute Myeloid Leukemias (AML) are a heterogeneous group of severe hematological malignancies resulting from the transformation of hematopoietic stem- or progenitor cells. Leukemic cells are blocked in the differentiation process, proliferate, invade the bone marrow and, thereby, disrupt normal hematopoiesis. AML primarily affect the elderly, with a median age at diagnosis around 70. Prognosis is poor and has not significantly improved in the past four decades, in particular for old patients (>60 years), their 5-year survival rate being close to 10%^1^. For fit patients, the standard treatment generally relies on intensive chemotherapy combining 3 days of an anthracycline (daunorubicine-DNR or idarubicine-IDA) and 7 days of cytarabine (Ara-C) (“3+7” regimen)^2^. Resistance to chemotherapy remains the main cause of relapse and death.

As compared to other cancers, clonal complexity of AML is relatively low at diagnosis^3^. However, standard AML chemotherapy can favor the emergence of chemoresistant clones through the elimination of those that are sensitive to the drugs (branched evolution), but also induce additional mutations entailing chemoresistance in the original predominant clone (linear evolution)^4^. The mechanisms responsible for chemoresistance acquisition and maintenance are not yet fully elucidated. Among those described are changes in the drug uptake/efflux^5^, modulation of the pro-drug activation process^6,7^, higher resistance to apoptosis^8^, enhanced DNA repair abilities^9^ and modulations of energetic and redox metabolisms^10,11^. A better characterization of the pathways dysregulated in chemoresistant AML would be instrumental to find new therapeutic targets to overcome chemoresistance. This could also provide new biomarkers allowing improvement of AML risk stratification, which is currently mostly based on the number and nature of the cytogenetic abnormalities^2^. Such prognosis biomarkers might help clinicians in their therapeutic decision, in particular for patients presenting high risks of therapy-related toxicity. This is all the more to be considered as new therapies, such as those targeting mutated FLT3 and IDH1/2 or BCL2, are emerging as promising alternative to standard chemotherapies^12^.

Here, we have approached the study of mechanisms underlying AML resistance through the identification of genes whose expression is dysregulated in AML cells resistant to DNR or Ara-C. We found that chemoresistant AML cells activate transcriptional programs implying the inflammatory response. In particular, we identified a strong up-regulation of the different subunits of the NADPH oxidase NOX2 and a large increase in ROS production in chemoresistant cells, particularly in those resistant to DNR. This activation of NOX2 participates in the acquisition/maintenance of the resistant phenotype. In addition, developing a score based on NOX2 subunits genes expression, we found that increased NOX2 level is a marker of both chemoresistance and adverse prognosis in AML patients.

## Materials and Methods

### Pharmacological inhibitors and reagents

Cytosine-β-D-arabinofuranoside (Ara-C), daunorubicin-hydrochloride (DNR) and Phorbol 12-myristate-13-acetate (PMA) were purchased from Sigma-Aldrich. The NADPH oxidase inhibitor VAS2870 was from Enzo Life Sciences. The luminescent ROS indicator L-012 was from Sobioda (ref W1W120-04891).

### Cell culture

HL-60 and U937 cells resistant to Ara-C and DNR were generated by culturing them in the presence of increasing concentrations of drugs as previously described^13^ and cloned using an Aria IIU cell sorter (Becton Dickinson).

### Patient samples

Bone marrow aspirates were collected after obtaining written informed consent from patients after approval by the institutional “Sud Méditerranée 1” Ethical Committee (ref 2013-A00260-45; HemoDiag collection). Immediately after collection, leukocytes were purified by density-based centrifugation using Histopaque 1077 (Sigma-Aldrich) and used for flow cytometry analysis. Clinical characteristics of the patients are provided in Supplementary Table 3. When indicated, cells were sorted after CD45/SSC gating^14^ with a CD45-Pacific-Blue antibody (Beckman Coulter) and CD34-PerCP-Vio700 (Miltenyi Biotec) using an Aria IIU cell sorter (Becton Dickinson).

### Flow cytometry

Cells were washed in PBS containing 2% FBS and incubated at 4°C for 30 minutes in the presence of FITC-conjugated gp91^phox^ antibodies (D162-4; MBL), washed and analyzed using LSR Fortessa flow cytometer (Becton Dickinson). For patient samples, leukemic cells were identified using CD45/SSC gating^14^ (see Supplementary Figure 3 for a gating example). The median fluorescence intensities for gp91^phox^ on the red blood cells present in the samples (negative control) was subtracted from the mean fluorescence intensities for gp91^phox^ on the leukemic cells.

### CRISPR/Cas9 knock-out of CYBB

A control or a *CYBB*-binding Cr-RNA (sequence available on request) was transfected together with a recombinant GFP-Cas9 protein and TracrRNA ATTO 550 (IDT) in HL-60 cells using the Amaxa technology. Twelve hours after transfection, GFP-positive cells were sorted and cloned. The DNA sequence surrounding the targeted CYBB sequence was PCR-amplified on genomic DNA from the selected clones, cloned in the TOPO-TA vector (Life Technologies) and sequenced.

### Cell proliferation and IC_50_ measurement

Cells were seeded at a concentration of 3×10^5^/mL in RPMI medium on day 0 and the number of cells was measured every two-three days using the MTS assay from Promega following the manufacturer’s protocol. For IC_50_ measurements, medium was complemented with increasing doses of DNR (Sigma-Aldrich). Viability was measured 24 hours later using the MTS assay. IC_50_ were calculated using the GraphPad PRISM software.

### RNA-seq libraries preparation and sequencing

Total RNAs were purified using the GenElute Mammalian Total RNA kit (Sigma-Aldrich), treated with DNase I (4U, New England Biolabs) and RNasin (2,5U, Promega) and re-purified. RNA quality was assessed using a BioAnalyzer Nano 6000 chip (Agilent). Three independent experiments were performed for each cell line. Libraries were prepared using TruSeq®Stranded mRNA Sample Preparation kit (Illumina). After the PCR amplification, PCR products were purified using AMPure XP beads (Agencourt Biosciences Corporation). The quality, size and concentration of cDNA libraries were checked using the Standard Sensitivity NGS kit Fragment Analyzer and qPCR (ROCHE Light Cycler 480). Libraries were sequenced using an Illumina Hiseq 2500 sequencer as single-end 125 base reads. Image analysis and base calling were performed using HiSeq Control Software (HCS), Real-Time Analysis (RTA) and bcl2fastq.

### RNA-seq mapping, quantification and differential analysis

RNA-seq reads were mapped on the Human reference genome (hg19, GRCh37p13) using TopHat2 (2.1.1)^15^ based on the Bowtie2 (2.3.5.1) aligner^16^. The reproducibility of replicates was verified using cufflinks v2.2.1 tool^17^ with the linear regression of reads per kilobase per million mapped reads (RPKM) between 2 replicates. Reads association with annotated gene regions was done with the HTseq-count tool v0.11.1^18^. Differential expression analysis was performed with DESeq2^19^ using normalization by sequencing depth and parametric negative binomial law to estimate the data dispersion. Genes with a fold change ≥2 or ≤ 0.5 and an adjusted p-value (FDR) < 0.05 were considered as differentially expressed genes (DEGs).

### GSEA, Gene Ontology and co-expression analysis

Ontology analyses were performed using the Panther software^20^ (http://www.pantherdb.org/). Gene Set Enrichment Analyses were performed using https://www.gsea-msigdb.org/gsea/index.jsp (version 4.0.3)^21^. Coexpression analyses were performed using the UCSC Xena browser (https://xenabrowser.net/heatmap/) with dataset from AML patients publicly available from the Cancer Genome Atlas Program (TCGA)^22^.

### RT-qPCR assays

Total mRNA was purified using the GenElute Mammalian Total RNA kit (Sigma-Aldrich). After DNase I treatment, 1 µg of total RNA was used for cDNA synthesis using the Maxima First Strand cDNA kit (ThermoFisher Scientific). qPCR assays were conducted using Taq platinum (Invitrogen) and the LightCycler 480 device (Roche) with specific DNA primers (IDT, sequence available on request). Data were normalized to the housekeeping *GAPDH* mRNA levels.

### NOX activity measurement

5×10^5^ cells were washed in PBS, resuspended in 100μl of PBS containing 10 mM glucose and 1mM CaCl_2_ prewarmed at 37°C and transferred to a white 96-well plate. PMA (100 nM) and VAS2870 (10 μM) were then added to the wells. The luminescent ROS indicator L-012 was then added at a final concentration of 500 μM and luminescence was recorded on a Centro XS^3^ LB 960 luminometer (Berthold Technologies).

### Generation of a prognosis score based on the expression of NOX2 genes

Gene expression microarray data from two independent cohorts of adult patients diagnosed with AML were used^23,24^. Patients without available survival data or with myelodysplastic syndrome diagnosis were excluded. The first cohort (Verhaak cohort)^23^ included 504 patients and the second cohort (Metzeler cohort)^24^ comprised 242 patients with normal karyotype (CN-AML). Affymetrix gene expression data are publicly available *via* the online Gene Expression Omnibus (http://www.ncbi.nlm.nih.gov/geo/) under accession numbers GSE6891 and GSE12417. *FLT3/NPM1* statuses for the Metzeler cohort patients were kindly provided by Metzeler et *al* ^24^. Because data normalization was different, expression level was normalized in both cohorts into standard normal cumulative distribution (X’ = (X-μ)/σ) for each gene. In the first cohort, cutpoints were determined for each probeset of interest using MaxStat analysis, therefore defining beta coefficients. The first cohort was then randomly divided in two sub-cohorts of 252 patients each (training and test sets). A gene expression-based risk score was created based on the levels of expression of NOX2 constituent genes in the training set. It was defined for each patient as the sum of the beta coefficients for all these genes weighted by +1 or −1 according to the patient sample signal above or below/equal the probeset MaxStat value as previously reported^25,26^. Patients from the training cohort were ranked according to increased NOX score and, for a given score value Y, the difference in survival of patients with a NOX score ≤Y or >Y was computed using MaxStat analysis. The NOX score was then individually computed for patients in the test set, using the cutoff value determined in the training cohort. Finally, we transposed our model in the validation cohort (Metzeler cohort), using both cutpoints defined for each probeset and the cutoff score determined in the training set. Survival analyses were assessed using Kaplan-Meier method and survival curves were compared using log-rank test. All statistical analyses were performed using the R.3.6.0 software.

### Statistical analyses

Results are presented as means ± S.D. Statistical analyses were performed using the paired Student’s t-test with the Prism 5 software. Differences were considered as significant for P-values of <0.05. *; **; *** correspond to P < 0.05; P < 0.01; P < 0.001, respectively. ns=not significant. Statistical analyses for the RNA-Seq experiments, GSEA and NOX score are described in the relevant sections.

## Results

### Chemoresistant AML cells activate inflammation-related transcriptional programs

To study AML chemoresistance, we performed RNA-Seq in the AML model cell line HL-60, using parental cells and DNR-resistant (DNR-R) or Ara-C-resistant (ARA-R) cell populations^13^. We identified 989 differentially expressed genes between drug-resistant and parental HL-60 cells (Figures 1A and 1B and Supplementary Table 1). Among them, some were up-regulated in resistant cells (452 for DNR-R and 283 for ARA-R) and others were down-regulated (312 for DNR-R and 146 for ARA-R). 132 genes were commonly up-regulated and 38 down-regulated in both types of resistant cells compared to parental cells (Figure 1A, 1B). In general, the most deregulated genes showed the same trend of up- or down-expression in both DNR-R and ARA-R cells (Figures 1C and 1D). However, many genes were preferentially modulated upon acquisition of resistance to one of the two drugs (594/764 for DNR-R and 259/429 for ARA-R) (Figures 1A and 1B and Supplementary Figure 1A). Ontological analysis of the genes up-regulated in both ARA-R and DNR-R cells showed a strong enrichment for immune signaling and inflammation (Figure 1E). An enrichment of few pathways involved in the regulation of transcription was found for the genes down-regulated in DNR-R cells, and no enrichment for specific pathways was found for the genes down-regulated in ARA-R cells (Supplementary Table 2). Gene set enrichment analysis (GSEA) further showed an enrichment of an inflammatory signature for the genes up-regulated in resistant *versus* parental HL-60 cells (Figure 1F). This signature included various cytokines and chemokines, which are up-regulated in both types of resistant cells, with, however, stronger induction in DNR-R cells (Supplementary Figure 1B). We then wondered if the inflammatory signature could also be enriched in patients relapsing after chemotherapy. To this end, we used published transcriptomic data obtained from 9 patients at diagnosis and relapse^27^. A GSEA analysis revealed that the inflammatory signature was enriched at relapse, specifically in the 3 patients from the FAB M2 subtype (Figure 1G). Accordingly, a signature containing all genes up-regulated in both ARA-R and DNR-R HL-60 cells was found enriched at relapse in these patients (Supplementary Figure 1C). Finally, the genes up-regulated specifically in DNR-R cells were also enriched in genes involved in inflammation whilst those enriched only in ARA-R cells were not enriched for any specific pathway (Supplementary Table 2). Thus, our data suggest that AML resistance to chemotherapies, in particular DNR, is associated with transcriptional reprogramming involving the induction of an inflammatory response.

**Figure 1:**
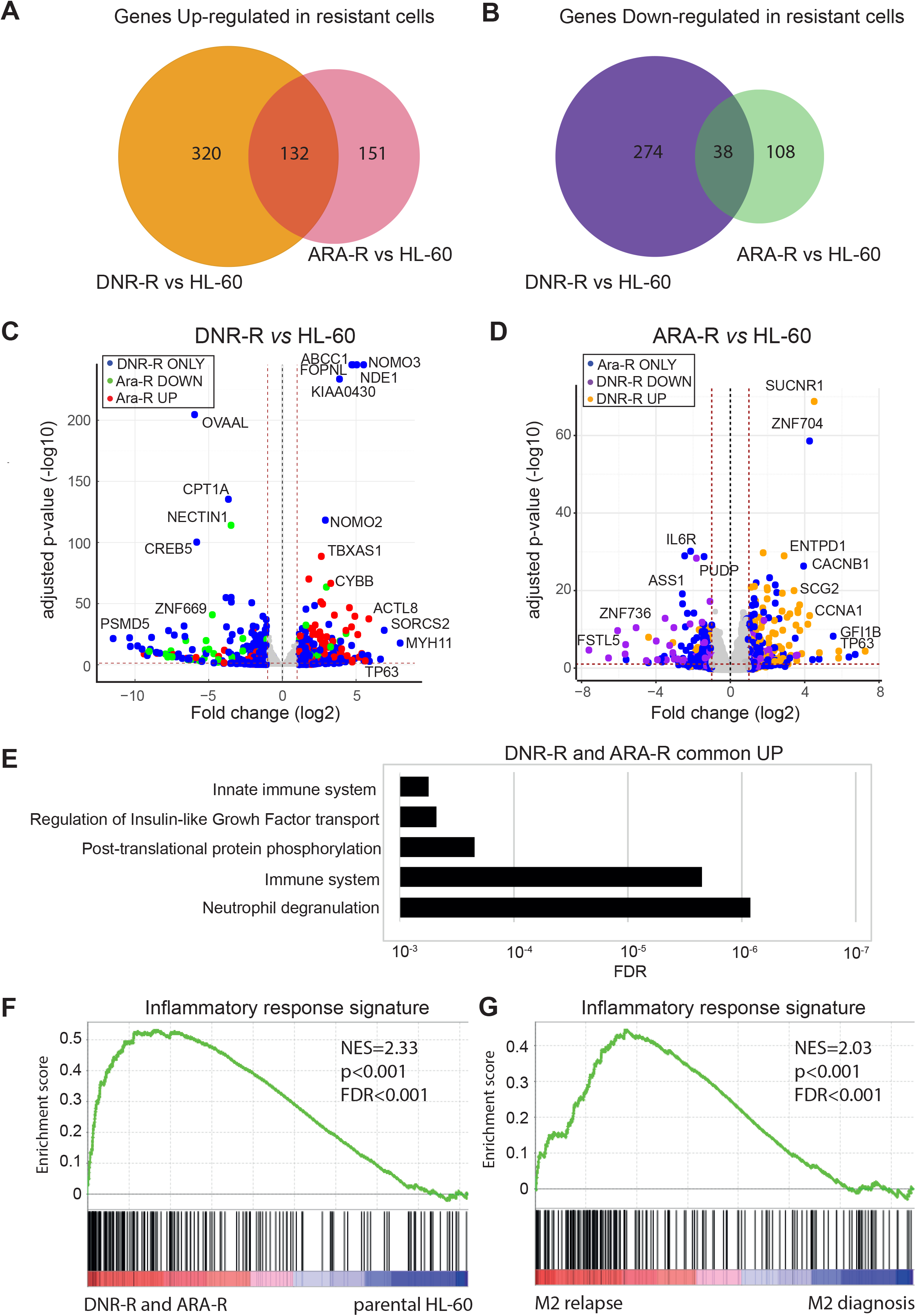
Transcriptional reprogramming upon acquisition of chemoresistance by HL-60 AML cells. RNA-Seq were performed on mRNAs purified in 3 independent experiments conducted with HL-60 cells and their Ara-C-resistant (ARA-R) and DNR-resistant (DNR-R) population derivatives. (A,B) Venn diagram for (A) up-regulated (>2 fold, FDR<0.05) and (B) down-regulated (>2 fold, FDR<0.05) genes in ARA-R and DNR-R cells compared to parental HL-60 cells. (C) Volcano plot for Differentially Expressed Genes (DEGs) between DNR-R- and HL-60 cells. Blue dots correspond to the DEGs in DNR-R but not in ARA-R cells, red dots to DEGs in DNR-R cells that are up-regulated in ARA-R cells and green dots to DEGs that are down-regulated in ARA-R cells. (D) Volcano plot for DEGs between ARA-R- and HL-60 cells. Blue dots correspond to the DEGs in ARA-R but not in DNR-R cells, yellow dots to DEGs in ARA-R cells that are up-regulated in DNR-R cells and purple dots to DEGs in ARA-R cells that are down-regulated in DNR-R cells. (E) Gene Ontology analysis of genes up-regulated in both DNR-R and ARA-R cells compared to parental HL-60 cells. (F) Gene Set Enrichment Analysis (GSEA) was performed using RNA-Seq data from ARA-R, DNR-R- and parental HL-60 cells. The enrichment for the “inflammatory response” signature (175 genes) is shown. NES (normalized enrichment score), nominal p-value and FDR (false decision rate) are presented. (G) The inflammatory signature (175 genes) identified in (F) was used in GSEA analysis with RNA-Seq data from 3 patients from the FAB M2 subtype obtained from a publicly available cohort^27^.

### Resistance to DNR is associated with strong overexpression and activation of the NAPDH oxidase NOX2

At the top of the list of genes up-regulated in the inflammatory response signature of resistant cells, in particular those resistant to DNR, we identified *CYBB* (Supplementary Figure 1B). It encodes the gp91^phox^ protein, the catalytic subunit of NOX2, the NADPH oxidase (NOX) family member principally expressed in the hematopoietic system and in AML^28^. Two of the NOX2 subunits, gp91^phox^ and p22^phox^ are associated with the plasma membrane (Figure 2A). Upon activation, the cytosolic subunits (p67^phox^, p47^phox^ and p40^phox^) and the small GTPase Rac2 are recruited to the membrane and activate the oxidase^29^. Further pointing to a link between NOX2 and DNR resistance, a NADPH oxidase signature was found to be strongly enriched in DNR-R cells compared to parental cells (Figure 2B). Indeed, although *CYBB* was found up-regulated in both ARA-R and DNR-R cells, the other NOX2 subunits were specifically up-regulated in DNR-R as compared to parental HL-60 cells (Figure 2C). RT-qPCR analysis confirmed that *CYBB* is strongly up-regulated in DNR-R cells and, to a lower extent, in ARA-R cells, as compared to parental HL-60 cells (Figure 2D). The *NCF2* gene (encoding p67^phox^) was also overexpressed in DNR-R but not in ARA-R cells (Figure 2D). To assess whether higher gene expression translates into higher NADPH oxidase activity, we measured extracellularly-produced superoxides (·O_2_-). Basal NADPH oxidase activity is low under standard growth conditions because the cytosolic subunits are not bound to the membrane. We therefore treated cells with PMA, a well-known activator of NADPH oxidases. No superoxide production was detected in HL-60 cells, which are known to have a low/absent NADPH oxidase activity when not differenciated^30^. A strong ROS production was measured in PMA-treated DNR-R but not in ARA-R cells (Figure 2E). Importantly, similar results were obtained in another AML cell line, U937, rendered resistant to Ara-C or DNR (Figure 2F). We then analyzed individual DNR-R or ARA-R clones (14 clones in each case) derived from the HL-60 cell populations used so far. All DNR-R clones showed a strong increase in both *CYBB* (Figure 3A) and *NCF2* (Figure 3B) mRNA levels (above 10 to more than 1000-fold compared to HL-60 cells). Although the Ara-C-resistant population showed no significant increase in ROS production (Figure 2E), several clones showed an increase in *CYBB* and *NCF2* expression in comparison to parental HL-60 cells (Figure 3A and 3B). This increase was however limited compared to DNR-R cells. Accordingly, PMA-induced extracellular ROS production was increased in all DNR-R clones compared to parental HL-60 and, to lower extents, in a fraction of ARA-R clones (Figure 3C and Supplementary Figure 2). To exclude the possibility that increased NOX2 expression and activity is due to the cloning procedure, we cloned the parental HL-60 cells. None of the HL-60 clones showed an increase in NADPH oxidase-derived ROS production (Supplementary Figure 2). Finally, we measured gp91^phox^ membrane expression using flow cytometry of two high ROS-producing clones (15165 and 15176) and two low ROS-producing clones (15160 and 15169) from DNR-R cells. Overexpression of gp91^phox^ at the cell surface was found to be high only in the clones producing high levels of ROS (15165 and 15176) (Figure 3D). VAS2870, a pan-NADPH oxidase inhibitor with preferential activity against NOX2^30^, strongly reduced PMA-induced ROS production in DNR-R clones (Figure 3E), further demonstrating that increased ROS production in chemoresistant AML cells is due to NADPH oxidase activation.

**Figure 2:**
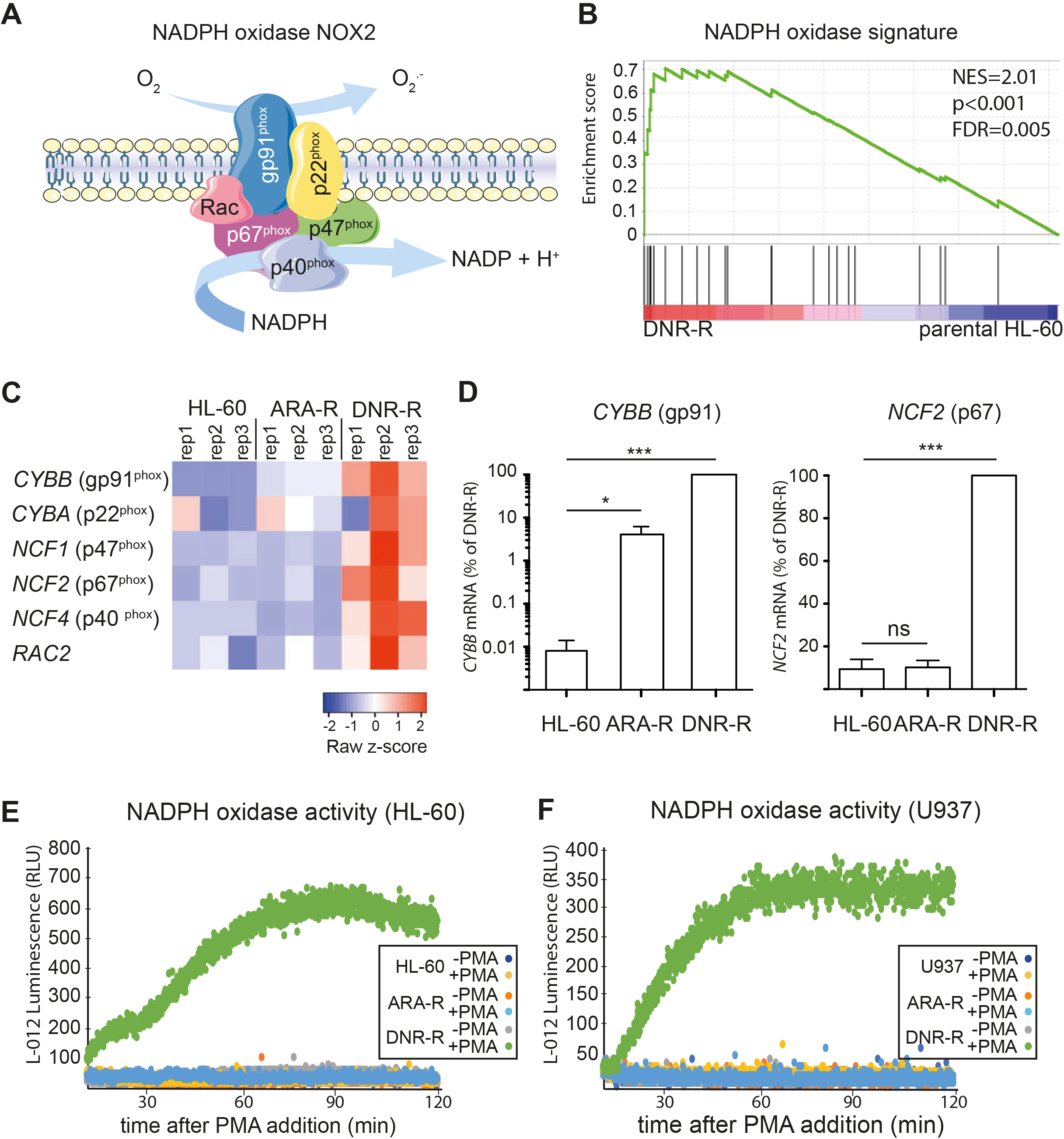
NOX2 expression and activity are increased in DNR-resistant HL-60 cells. (A) The NAPDH oxidase NOX2 is composed of two membrane-associated proteins, gp91^phox^ and p22^phox^. Upon activation, the cytosolic subunits p67^phox^, p47^phox^ and p40^phox^ translocate to the membrane together with the small GTPase Rac2 to form the full oxidase complex. (B) GSEA was performed using RNA-Seq data from DNR-R and parental HL-60 cells. The enrichment for the “NADPH oxidase” signature (23 genes) is shown. NES (normalized enrichment score), nominal p-value and FDR are presented. (C) Heatmap of the RNA-seq data in parental, DNR-R and ARA-R cells showing the expression of the genes encoding the NOX2 subunits (*CYBB*, *CYBA*, NCF1, *NCF2*, *NCF4*, *RAC2*). (D) The expression of *CYBB* and *NCF2* was analyzed by RT-qPCR on mRNA purified from parental-, ARA-R- and DNR-R cells and normalized to *GAPDH* mRNA levels. Data are presented as percentages of the DNR-R condition (n=5, mean +/− SD). (E,F) Parental, ARA-R and DNR-R HL-60 (A) or U937 (B) cells were treated or not with PMA and the production of ROS was measured by following L-012 luminescence over 2 hours (n=4 for HL-60, n=3 for U937, a representative experiment is shown).

**Figure 3:**
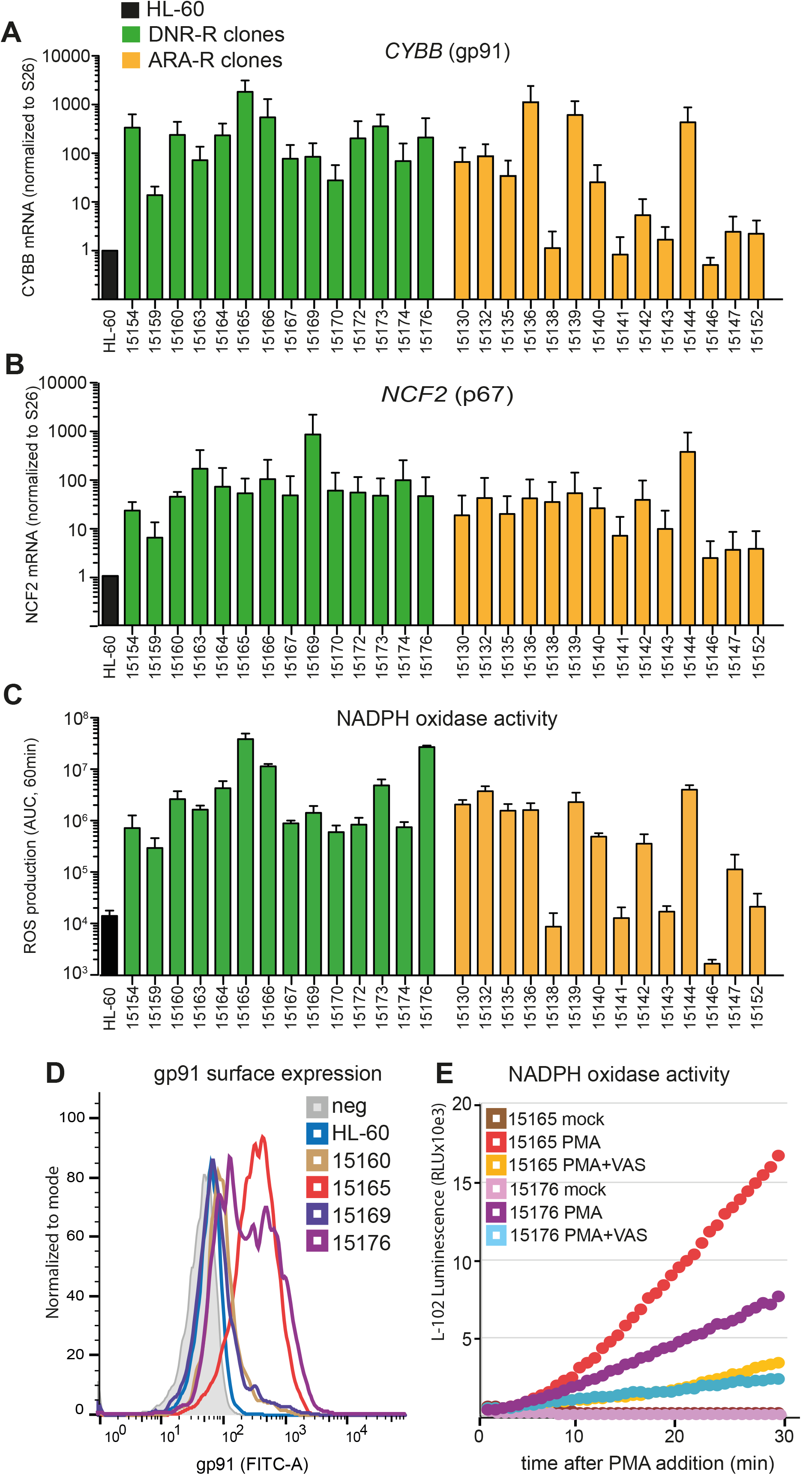
Overexpression of NADPH oxidase components is responsible for increased ROS production in chemoresistant AML cells. (A-C) DNR-R and ARA-R cell populations were cloned and 14 clones for each were used to measure mRNA expression for *CYBB* (A) and *NCF2* (B) using RT-qPCR. Data were normalized to *S26* mRNA levels and are presented as ratios to parental HL-60 cells (n=3, mean +/− SD). (C) ROS production was measured after addition of PMA using L-012 luminescence. The AUC (Area Under Curve) was calculated after plotting L-012 luminescence over 1 hr (n=3, mean +/− SD). (D) gp91^phox^ surface expression was measured by flow cytometry in parental HL-60 cells as well as in two DNR-R clones showing high NADPH oxidase activity (15165 and 15176) and two clones with low NADPH oxidase activity (15160 and 15169). (E) DNR-R clones 15165 and 15176 were stimulated or not with PMA with or without VAS2870 and ROS production was measured by luminometry (n=3, a representative experiment is shown).

Altogether, our data suggest that acquisition of chemoresistance, in particular to DNR, is associated with increased expression of the subunits of the NOX2 NADPH oxidase, its cell surface expression and activity in AML cells.

### gp91^phox^ overexpression participates in the resistance to DNR

Using the CRISPR/Cas9 technology, we knocked out the *CYBB* gene in the DNR-R clone showing the highest level of gp91^phox^ expression (clone 15165). This led to a strong decrease in gp91^phox^ cell surface expression (Figure 4A) and abolished NADPH oxidase activity (Figure 4B). No difference was observed between the proliferation rates of parental HL-60 and DNR-R cells expressing or not gp91^phox^ (Figure 4C). However, DNR-R cells CRISPRed for gp91^phox^ regained sensitivity to DNR compared to control CRISPRed cells (Figure 4D). This suggests that over-expression of gp91^phox^ contributes to the acquisition and/or the maintenance of resistance to DNR in our model.

**Figure 4:**
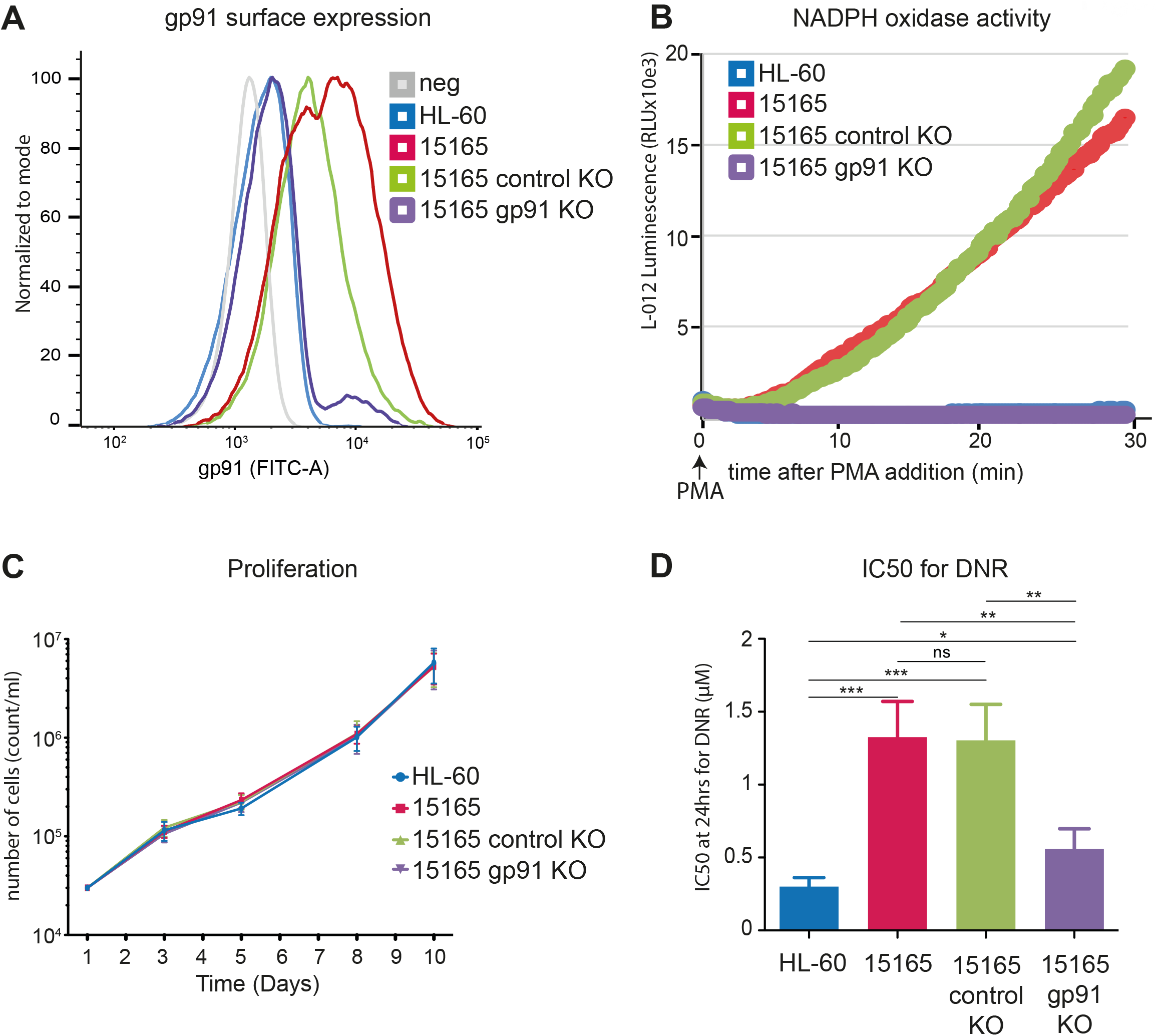
NOX2 overexpression participates in the acquisition/maintenance of resistance to DNR. (A) gp91^phox^ surface expression was measured by flow cytometry in parental HL-60 cells, in DNR-R clones (15165) as well as 15165 cells CRISPR/Cas9 KO for gp91^Phox^ and −15 165 cells subjected to mock CRISPR/Cas9 mutagenesis. (B) NADPH oxidase activity was detected by ROS measurement in the cells presented in (A) after PMA addition using L-012 luminescence. (n=3, a representative experiment is shown). (C) Cell proliferation was measured using MTS (n=3, mean +/− SD). (D) IC_50_ for DNR was measured after 24 hrs of treatment using MTS (n=4, mean +/− SD).

### gp91^phox^ expression and NOX2 activity is higher in FAB M4/M5 AML subtypes

To further characterize the importance of NOX2 in AML patients, we flow cytometry-assayed the expression of gp91^phox^ on the surface of primary AML cells from 74 patients at diagnosis. Differences in gp91^phox^ levels were neither associated with a specific group from the ELN-2017 classification^2^ (favorable, intermediate or adverse), nor with the NPM1 or FLT3 mutational status (Supplementary Figure 4 and Supplementary Table 3). However, gp91^phox^ levels were significantly higher in the M5 AML subtypes compared to normal CD34^+^ cells and leukemic blasts from the M0, M1 and M2 subtypes from the French-American British (FAB) classification (Figure 5A). Using transcriptomic data from the TCGA cohort^22^, we confirmed that *CYBB* mRNA expression is higher in the M4/M5 subtypes (Figure 5B). To assess whether increased gp91^phox^ cell surface expression is linked to higher NAPDH oxidase activity, we measured PMA-stimulated ROS production using sorted AML cells (bulk CD34^-^ leukemic cells and Leukemic Stem Cells (LSC)-containing CD34^+^) from one patient with low gp91^phox^ (M1 subtype, patient #17215) and another with a high gp91^phox^ cell surface expression (M4 subtype, patient #17226) (Figure 5C). Weak PMA-induced ROS production was detected in the low gp91^phox^-expressing patient cells contrasting with high production in high gp91^phox^-expressing patient cells (Figure 5D). For the M4 patient cells, gp91^phox^ level (Figure 5C) and ROS production (Figure 5D) were much higher in the bulk of leukemic cells (CD34^-^) than in the LSC-containing compartment (CD34^+^).

**Figure 5:**
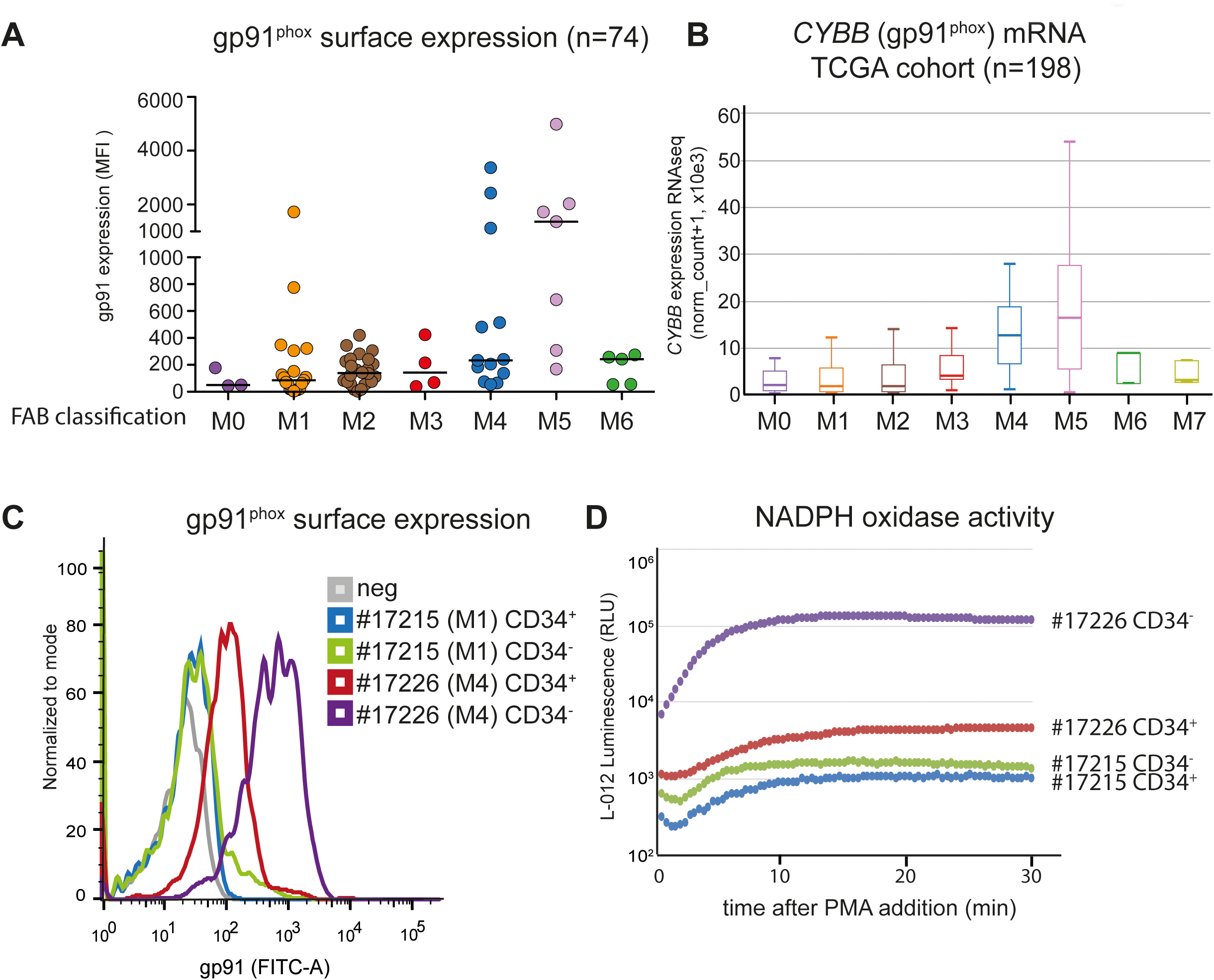
NOX2 expression and activity is higher in FAB M4/M5 patients. (A) gp91^phox^ expression (MFI) was measured by flow cytometry on the leukemic cells (CD45/SSC gating) present in bone marrow aspirates taken at diagnosis from 74 AML patients. Patients are classified according to the FAB classification. The median is represented as a horizontal line for each group. p=0.0057 in one-way Anova (Kruskal-Wallis test) (B) mRNA expression for *CYBB* gene was measured by RNA-Seq in the TCGA cohort comprising 198 AML patients. (C) gp91^phox^ surface expression was measured by flow cytometry on AML cells from two patients sorted using CD45/SSC gating and separated according to their level of expression of the CD34 marker. (D) Sorted AML cells (CD34^+^ and CD34^-^) for each patient were used to measure ROS production after stimulation with PMA (10 nM).

### NOX2 overexpression is a marker of adverse prognosis in AML

To study the link between NOX2 overexpression and AML response to treatments, we analyzed the prognosis value of the *CYBB* and *NCF2* genes in two independent cohorts of AML patients^23,24^. Their overexpression at diagnosis was associated with a significantly lower patient survival (Figure 6A). We then determined the individual prognosis value of the genes coding for the six NOX2 subunits (*CYBB*, *CYBA*, *NCF1*, *NCF2*, *NCF4*, *RAC2*) in the Verhaak cohort. Increased expression was associated with a bad prognosis for four of them (*CYBB*, *NCF2*, *NCF4*, *RAC2*) and with a good prognosis for the other two (*CYBA* and *NCF1*) (Table 1). To take into account the expression of all NOX2 subunits instead of individual ones, we developed a NOX2 subunits gene expression-based score (see methods). The contribution of individual NOX2 genes and the cutpoint were determined using a training cohort comprising 252 patients (Verhaak training set)^23^. Patients with a high “NOX score” had a worse prognosis with a hazard ratio of 1.85 in univariate Cox analysis (Figure 6B and Table 2). The prognosis significance of the NOX score was then confirmed in a test set (n=252) from the Verhaak cohort (Figure 6C and Table 3) as well as an independent validation cohort (Metzeler cohort^24^, n=240)(Figure 6C and Table 4). Finally, the poor prognostic-based NOX score remained statistically significant in a multivariate Cox analysis including the cytogenetic-based classification, the NPM1/FLT3 mutational status (for the normal karyotype cohort) and age (Table 2–4). To determine if the worse prognosis of high NOX score patients could be associated to higher chemoresistance, we selected the 10% of patients with the highest NOX score and compared their transcriptome to those of the 50% of patients with the lowest NOX scores. We then used the 50 most up-regulated genes in the high NOX score patients (Supplementary Table 4) as a gene-set in a GSEA analysis of the RNA-Seq data from parental and chemoresistant HL-60 cells. The high NOX score gene set was significantly enriched in the chemoresistant compared to parental HL-60 cells (Figure 6D). Interestingly, the most up-regulated gene in the high NOX score patients is *NAMPT*, encoding a critical enzyme in the recycling of NAD, an essential co-factor for NADPH oxidase activity. Finally, *NAMPT* expression is highly correlated to that of the *CYBB*, *NCF1* and *NCF2* genes in AML patients (Figure 6E).

**Figure 6:**
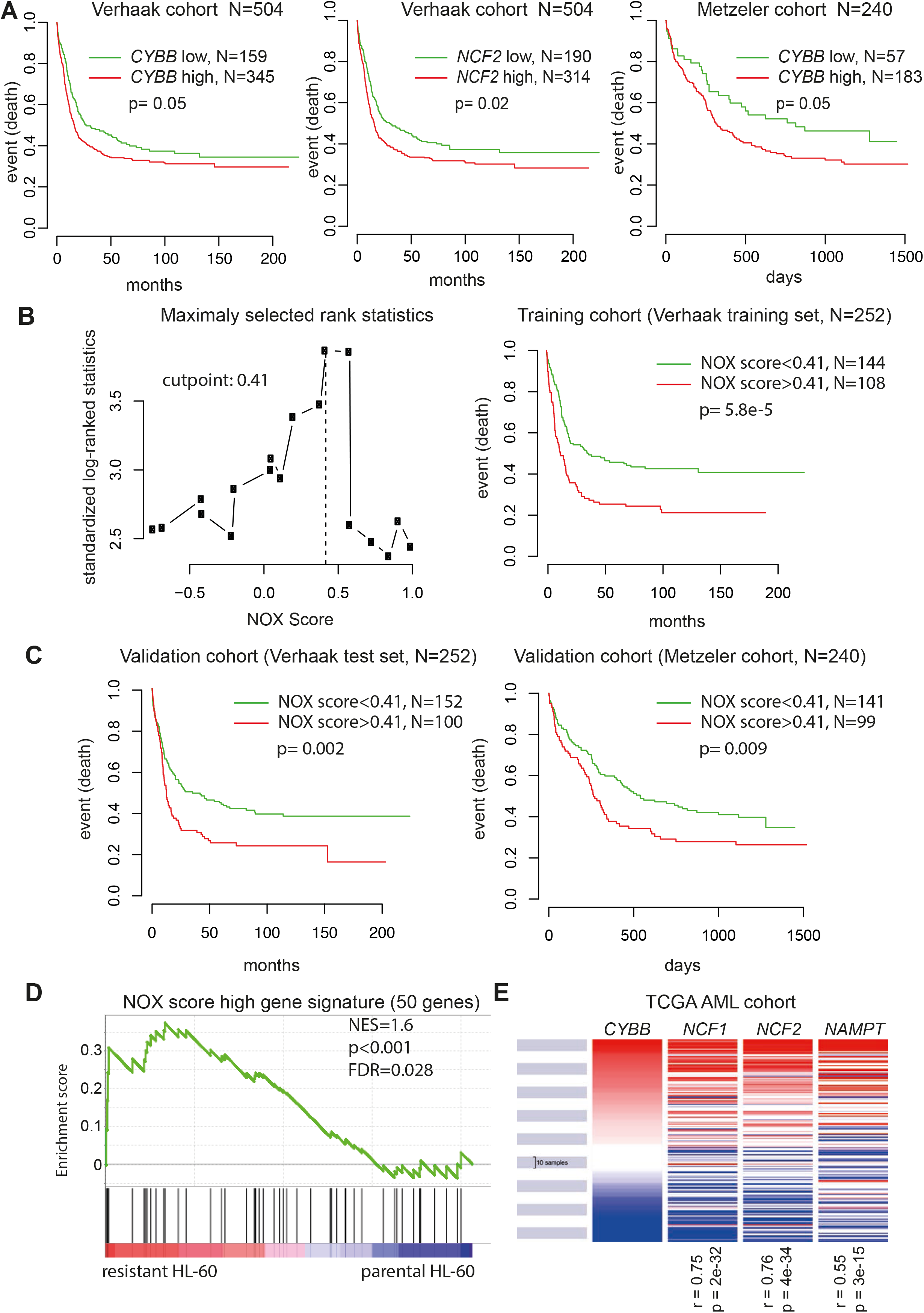
Identification of a NOX score with prognosis value in AML. (A) Kaplan-Meyer survival curves according to the levels of expression of *CYBB* and *NCF2* in two independent cohorts. p-values are determined with log-ranked test. (B) Patients of the training cohort comprising 252 patients were ranked according to increasing NOX score. The cut-point of 0.41 was selected using the MaxStat R function to obtain a maximum difference in overall survival (OS) between the two groups. Kaplan-Meyer survival curves were established using the cut-point value of 0.41 in the training cohort. (C) The NOX-score was validated in two independent cohorts using the 0.41 cutpoint. (D) Gene Set Enrichment Analysis (GSEA) was performed using RNA-Seq data from chemoresistant (ARA-R and DNR-R) compared to parental HL-60 cells. The gene list comprises the 50 most up-regulated genes in the 10% of patients with the highest NOX score, compared to the 50% patients with the lowest NOX score in the Verhaak cohort. NES (normalized enrichment score) and nominal p-value are presented. (E) Expression of *CYBB*, *NCF1*, *NCF2* and *NAMPT* in 200 patient samples from the TCGA cohort. R represents the Pearson rho correlation coefficient between the expression of *NCF1*, *NCF2* or *NAMPT* and *CYBB*.

**Table 1.** List of the 6 probesets included in the NOX score. *Gene symbol, adjusted p-value, hazard ratio and prognosis significance are provided for each gene, as determined in the Verhaak cohort (n=504)*.

| Probe set | Gene symbol | Benjamini Hochberg corrected p-value | Hazard ratio | Prognosis |
| --- | --- | --- | --- | --- |
| 229445_at | CYBA | 0.0104 | 0.673 | Good |
| 203922_s_at | CYBB | 0.0511 | 1.263 | Bad |
| 204961_s_at | NCF1 | 0.0172 | 0.677 | Good |
| 209949_at | NCF2 | 0.0208 | 1.303 | Bad |
| 205147_x_at | NCF4 | 0.0169 | 1.359 | Bad |
| 207419_s_at | RAC2 | 0.0058 | 1.490 | Bad |

**Table 2.** Cox analysis of overall survival in the Verhaak training set (n=252) according to NOX score, cytogenetic prognosis, and age. *Hazard ratio (HR) and p-values are shown for each parameter in univariate and multivariate Cox analyses*.

| Scores | Univariate Cox analysis |  | Multivariate Cox analysis |  |
| --- | --- | --- | --- | --- |
|  | HR | p-value | HR | p-value |
| NOX score | 1.85 | 7.54e-05 | 1.76 | 1.48e-03 |
| Cytogenetic prognosis | 1.85 | 2.87e-06 | 1.63 | 3.32e-04 |
| Age (per year) | 1.02 | 8.23e-03 | 1.02 | 0.044 |

**Table 3.** Cox analysis of overall survival in Verhaak test set (n=252) according to Nox score, cytogenetic prognosis, and age. *Hazard ratio (HR) and p-values are shown for each parameter in univariate and multivariate Cox analyses*.

| Scores | Univariate Cox analysis |  | Multivariate Cox analysis |  |
| --- | --- | --- | --- | --- |
|  | HR | p-value | HR | p-value |
| NOX score | 1.59 | 2.91e-03 | 1.54 | 0.014 |
| Cytogenetic prognosis | 1.72 | 1.59e-04 | 1.65 | 4.05e-04 |
| Age (per year) | 1.01 | 0.095 | not included |  |

**Table 4.**
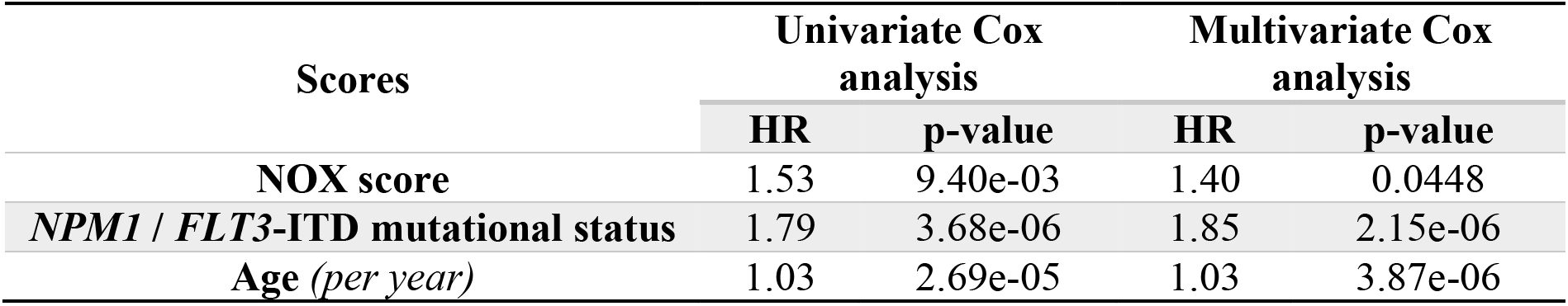
Cox analysis of overall survival in the Metzeler validation set (n=240) according to NOX score, *NPM1* & *FLT3* mutational status, and age. *Hazard ratio (HR) and p-values are shown for each parameter in univariate and multivariate Cox analyses. ITD: internal tandem duplication*.

Altogether, this suggested that overexpression of the NOX2 NADPH oxidase is a marker of adverse prognosis in AML associated with enhanced chemoresistance of the leukemic cells.

## Discussion

Our work unveiled that NOX2 expression increases in chemoresistant AML and could participate to the acquisition/maintenance of chemoresistance. Moreover, NOX2 was found to be a marker of adverse prognosis in AML patients. NOX2 is the main NADPH oxidase complex expressed in the hematopoietic system, in particular in monocytes/macrophages. It is responsible for the respiratory burst, a release of superoxides (·O_2_-) in the phagosome, which is used to eliminate engulfed pathogens, in particular through its transformation to hypochlorous acid. Mutations in the six NOX2 subunits were found to cause Chronic Granulomatous Disease, a genetic disease leading to severe immunodeficiency and life-threatening infections^33^. In addition to their essential role in host protection, ROS produced by NADPH oxidases also function as second messengers in the regulation of numerous signaling pathways^34^. Dysregulations of NADPH oxidases expression and activity have been linked to various cancers, including AML^35^. NOX2 expression was found to be generally higher in leukemic blasts from AML patients compared to normal CD34^+^ hematopoietic progenitors and involved in leukemic cells proliferation^36^. NOX2 was shown to be mostly expressed in M4 and M5 subtypes of the FAB classification and rarely on leukemic blasts from M1 or M2 subtypes^37^. By analyzing the gp91^phox^, the catalytic subunit of NOX2, at the surface of blasts from 74 AML patients, we confirmed that NOX2 expression is higher in the M4/M5 subtypes than in the M0, M1 and M2 subtypes. In addition, within a given patient sample, gp91^phox^ expression and NOX2 activity were found higher in the bulk of leukemic cells (CD34^-^) compared to the LSC-containing CD34^+^ compartment, which nevertheless showed basal activity. This is coherent with the fact that NOX2 is required for LSC self-renewal and, in turn, leukemogenesis^28^. Moreover, our results indicate that the increase in NOX2 activity in chemoresistant cell lines is linked with the transcriptional activation of most, if not all, of its six subunits. Interestingly, their expression, in particular those of *CYBB*, *NCF1* and *NCF2*, is highly correlated. Specific transcription factors that govern their expression could thus be up-regulated upon acquisition of chemoresistance. PU-1 is the main transcription factor involved in the transcriptional regulation of both *CYBB* and *NCF2* in myeloid cells^38,39^. We only detected minor changes in PU-1 expression between DNR-resistant and parental HL-60 cells (1.28-fold induction, p=0.09). Therefore, higher NOX2 expression in chemoresistant cells is unlikely to be due to increased PU-1 levels but rather to still-to-be-identified transcription factors and/or functional activation of PU-1.

Increased NOX2 expression could confer a selective advantage to the chemoresistant clones. We show here that preventing NOX2 activation in DNR-R cells through the deletion of *CYBB* re-sensitized them to DNR. This indicates that NOX2 overexpression could be involved in the acquisition and/or maintenance of resistance. The underlying molecular mechanisms remain to be characterized. They might involve the regulation of specific signaling pathways through the reversible oxidation of cysteines present in the catalytic site of enzymes such as kinases and phosphatases^40^ or, as we showed previously, of SUMOylation enzymes^41,42^. Their modulation would participate in the transcriptional reprogramming observed in chemoresistant AML cells and participate in the acquisition and/or maintenance of chemoresistance. NOX-derived ROS were also shown to promote AML cell proliferation^36,43^, in particular by activating genes involved in glycolysis^44^. In addition, oncogenes such as mutated *RAS*^45,46^ or *FLT3-ITD*^47^ activate NADPH-oxidase-derived ROS production, which increases hematopoietic cell proliferation and leukemic transformation. Finally, NOX2-derived ROS stimulate the transfer of mitochondria from bone marrow stromal cells to the AML blasts, increasing their metabolic activity and proliferation^48^. However, in our study, high NOX-expressing chemoresistant clones did not proliferate faster than chemosensitive parental cells and deletion of *CYBB* did not affect their doubling time. Thus, although the pro-proliferative effect of NOX-derived ROS is involved in leukemogenesis, it does not seem to confer a proliferative advantage to chemoresistant cells. Finally, NADPH-oxidase derived ROS are also involved in the interaction between leukemic cells and the immune system. They can induce apoptosis of Natural Killers and T-cells^37^ and prevent the maturation of Dendritic Cells^49^. Whether such an inactivation of anti-tumor immune cells could also confer a selective advantage to the chemoresistant AML cells that overexpress NOX2 remains an open question.

A selective advantage of high NOX2-expressing chemoresistant leukemic cells could explain that high NOX2 expression is a marker of adverse prognosis in AML. The NOX score we developed is independent from other prognosis factors such as age and cytogenetic risk or *NPM1/FLT3* mutational status in normal karyotype AML. It should now be validated in a prospective cohort to confirm its prognosis value. NAPDH-oxidase inhibitors have been proposed as anti-cancer drugs. Histamine, which inhibits NADPH-oxidase-derived ROS production, is already approved, in combination with IL-2, to prevent relapse in AML^50^. Many other molecules targeting NADPH oxidases have been developed in the past few years^51^. However, most of them suffer from a lack of specificity^52,53^. Interestingly, we found that NAMPT, a rate-limiting enzyme in the recycling of NAD, is the most upregulated gene in the high NOX score patients and its expression is highly correlated to NOX2 subunits levels in AML patients. In addition, NAMPT was more expressed in chemoresistant HL-60 cells than in parental cells, in line with recent findings showing its overexpression in chemoresistant *versus* chemosensitive LSC^54^. Inhibitors of NAMPT have shown anti-leukemic activity in preclinical models^54–56^ and some of them are being tested in clinical trials (clinicaltrials.gov identifiers NCT02702492, NCT04281420). Thus, targeting NAMPT in high NOX score-expressing patients could, by decreasing NADPH supply, be another way to inhibit NOX2 activity in AML cells. Altogether, our work point to a link between NOX2 overexpression and AML chemoresistance. Its targeting might constitute a new therapeutic strategy to overcome AML chemoresistance and improve their prognosis.

## Acknowledgments

We are grateful to the IGMM “Ubiquitin Family in Hematological Malignancies” group members for fruitful discussions and critical reading of the manuscript. We thank the Montpellier Ressources Imagerie (MRI) platforms for technical assistance. Funding was provided by the CNRS, Ligue Nationale contre le Cancer (Programme Equipe Labellisée), the INCA (ROSAML), the ANR (“Investissements d’avenir” program; ANR-16-IDEX-0006), the Fondation pour la Recherche Médicale (FRM) to LG, the Fondation ARC to PG and the Ligue Nationale contre le Cancer to MB. The HEMODIAG_2020 collection of clinical data and patient samples was funded by the Montpellier University Hospital, the Montpellier SIRIC and the Languedoc-Roussillon Region. MGX acknowledges financial support from the France Génomique National infrastructure, funded as part of “Investissements d’Avenir” program managed by the Agence Nationale pour la Recherche (contract ANR-10-INBS-09).

## Supplementary Material Legends

**Supplementary Figure 1: Genes activated in chemoresistant AML cell lines are linked with the inflammatory response and enriched in chemoresistant patients.** (A) Heatmap for the 50 most differentially expressed genes between parental-, DNR-R- and ARA-R cells. All biological replicates are shown. (B) Heatmap for the top 25 genes from the “inflammatory response” signature in DNR-R and parental HL-60 cells. (C) The list of genes induced in both DNR-R and ARA-R compared to parental HL-60 was used in GSEA analysis with RNA-Seq data from 3 patients from the FAB M2 subtype obtained from a publicly available cohort^27^.

**Supplementary Figure 2: Measure of NOX2 activity in parental and chemoresistant HL60 clones.** ROS production was measured after addition of PMA using L-012 luminescence in (A) DNR-R, (B) ARA-R and (C) HL-60 clones over 1 hr (n=3, a representative experiment is shown).

**Supplementary Figure 3: Gating strategy for patient samples.** Cells are first distinguished from debris and dead cells using FSC/SSC gating. Using CD45/SSC plot, the leukemic cells are distinguished from other populations present in the sample as CD45^dim^/SSC^low^. Erythrocytes are CD45^neg^/SSC^low^ while lymphocytes CD45^high^/SSC^low^.

**Supplementary Figure 4: gp91phox expression in AML cells is not correlated with ENL2017 risk classification, NPM1 or FLT3 mutational status.** gp91^phox^ expression (MFI) was measured by flow cytometry on bone marrow aspirates from 74 patients taken at diagnosis. Patients are classified according to (A) the ELN-2017 classification, (B) NPM1 mutational status or (C) FLT3 mutational status. No significant differences between groups using Mann-Whitney test.

**Supplementary Table 1: Differentially Expressed Genes between DNR-R and ARA-R cells compared to parental HL-60 cells.**

**Supplementary Table 2: Ontology analysis of up- and down-regulated genes in DNR-R and ARA-R compared to parental HL-60.**

**Supplementary Table 3: Clinical characteristics of the patient included in this study**

**Supplementary Table 4: High NOX score gene signature.** Mean expression for each Affimetrix probe for all patients from the Verhack cohort who had a NOX score above 1.37 (10% of patients) was divided by the mean expression for the patients who had a NOX score below 0.41 (50% of patients). The 50 genes having the highest ratio are presented.

