## Supplementary figures and images for "The NADPH oxidase NOX2 is a marker of adverse prognosis involved in chemoresistance of Acute Myeloid Leukemias"

### Supplementary Figures

#### Supplementary Figure 1

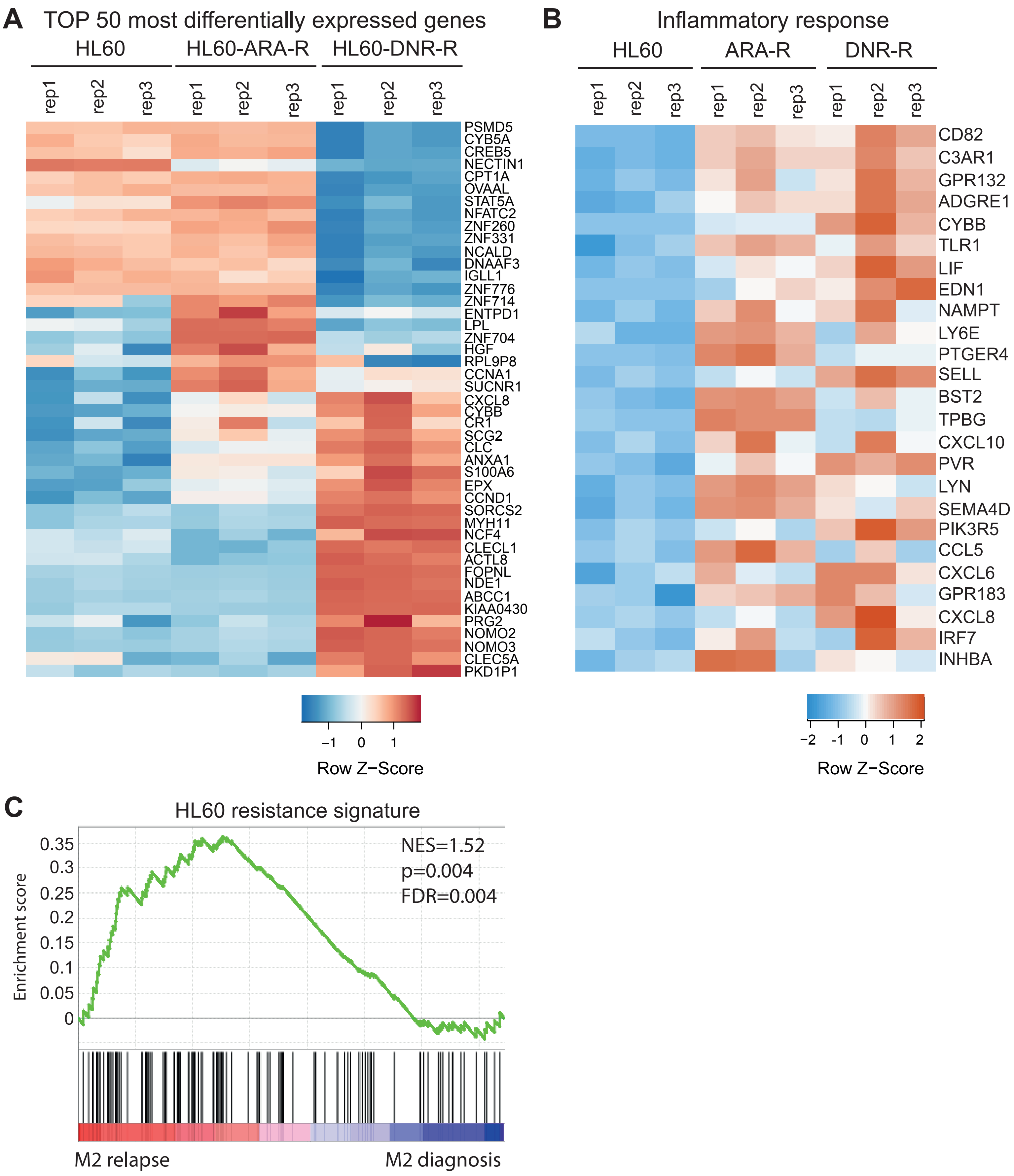

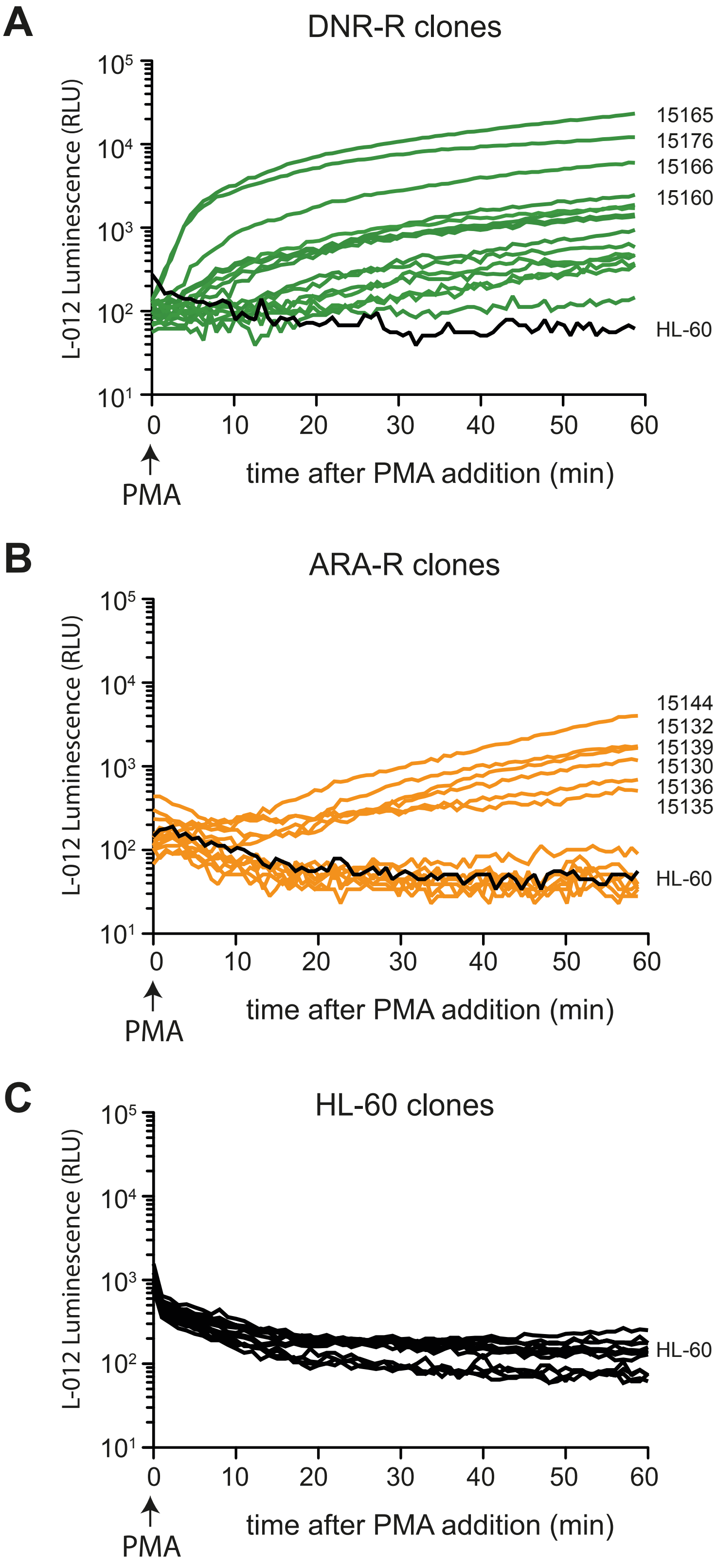

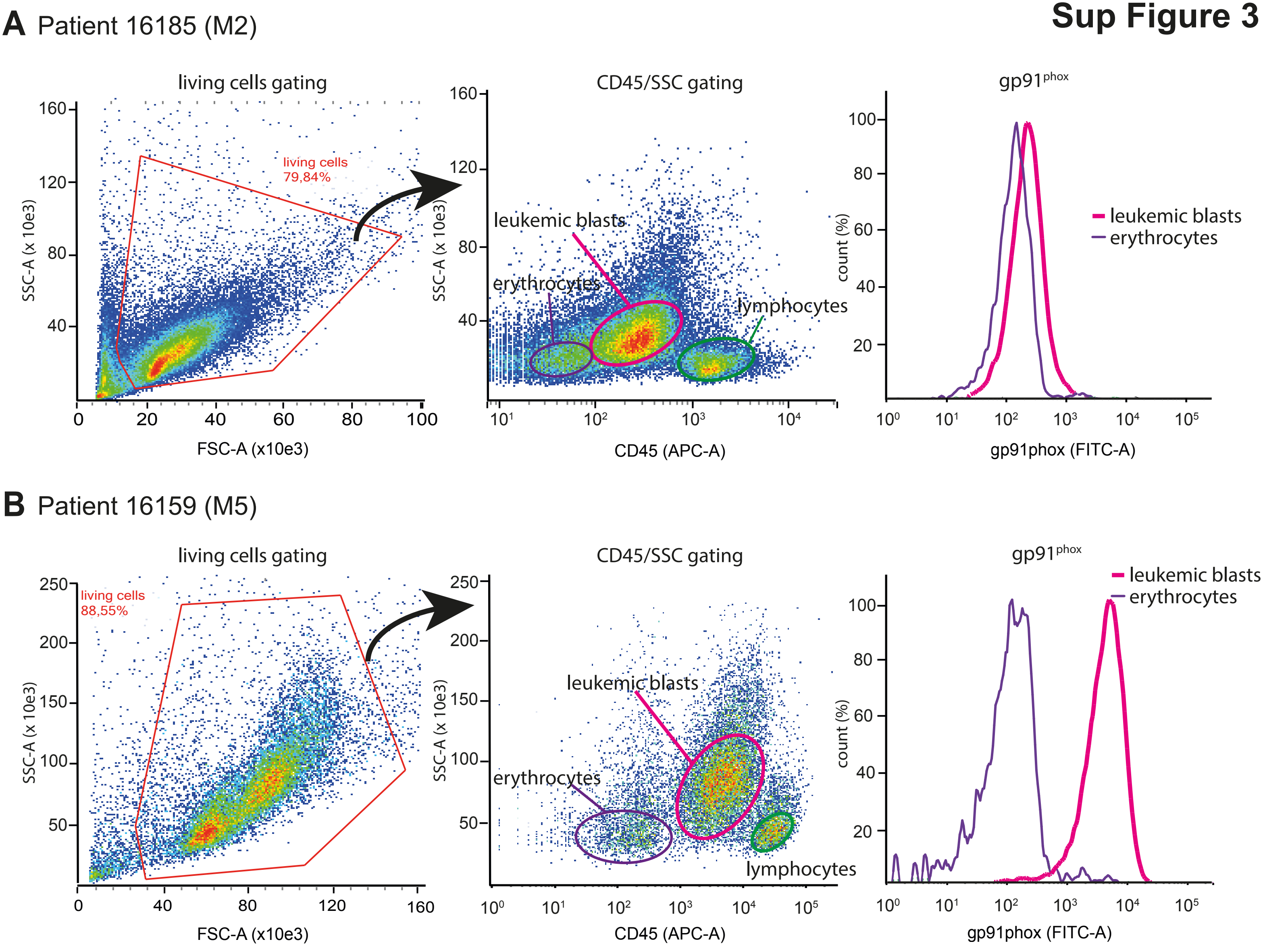

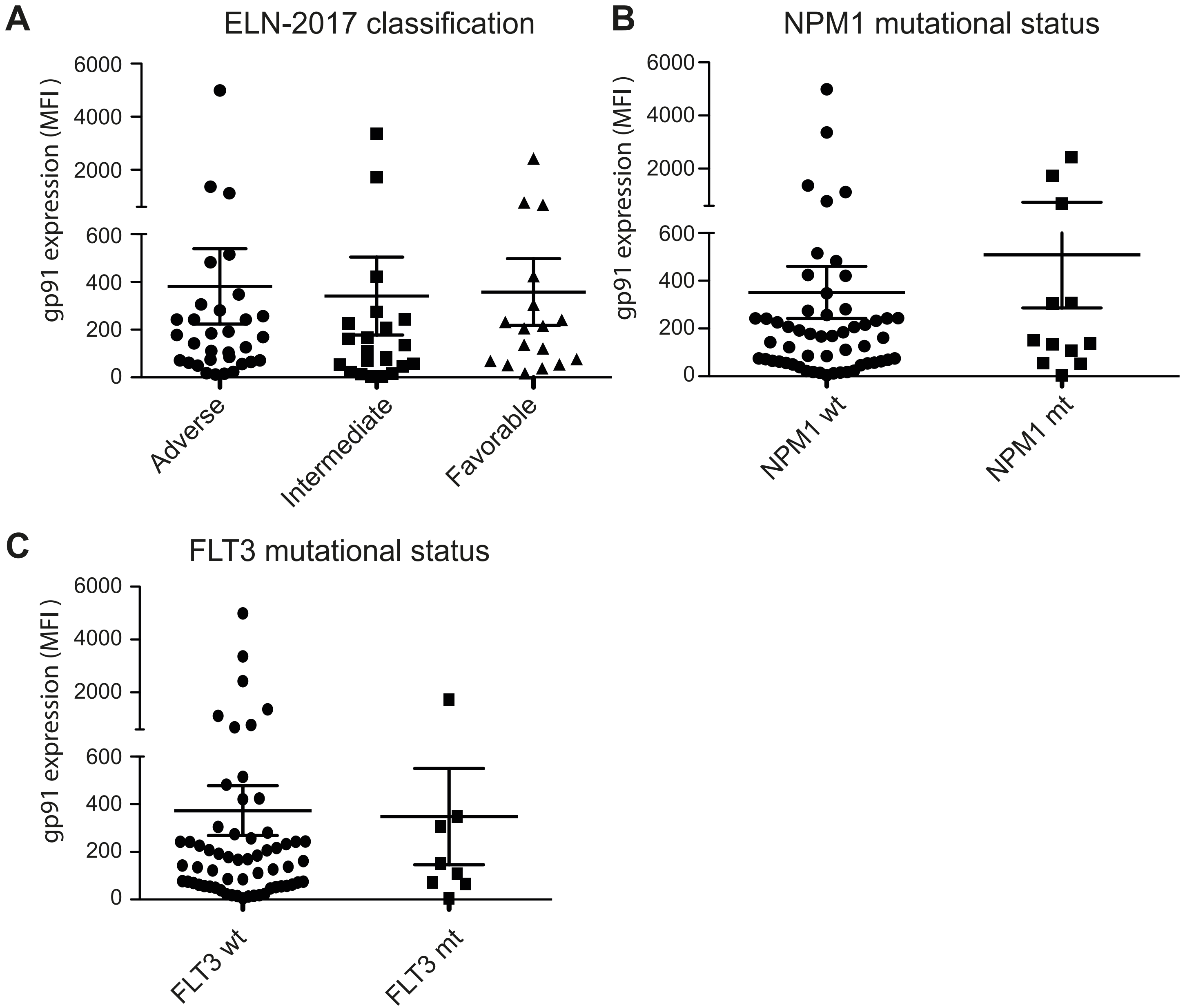
